# Integrated Structure-Transcription analysis of small molecules reveals widespread noise in drug-induced transcriptional responses and a transcriptional signature for drug-induced phospholipidosis

**DOI:** 10.1101/119990

**Authors:** Francesco Sirci, Francesco Napolitano, Sandra Pisonero-Vaquero, Diego Carrella, Diego L. Medina, Diego di Bernardo

## Abstract

We performed an integrated analysis of drug chemical structures and drug-induced transcriptional responses. We demonstrated that a network representing 3D structural similarities among 5,452 compounds can be used to automatically group together drugs with similar scaffolds and mode-of-action. We then compared the structural network to a network representing transcriptional similarities among a subset of 1,309 drugs for which transcriptional response were available in the Connectivity Map dataset. Analysis of structurally similar, but transcriptionally different, drugs sharing the same mode of action (MOA) enabled us to detect and remove weak and noisy transcriptional responses, greatly enhancing the reliability and usefulness of transcription-based approaches to drug discovery and drug repositioning. Analysis of transcriptionally similar, but structurally different drugs with unrelated MOA, led us to the identification of a “toxic” transcriptional signature indicative of lysosomal stress (lysosomotropism) and lipid accumulation (phospholipidosis) partially masking the target-specific transcriptional effects of these drugs. We further demonstrated by High Content Screening that this transcriptional signature is caused by the activation of the transcription factor TFEB, a master regulator of lysosomal biogenesis and autophagy. Our results show that chemical structures and transcriptional profiles provide complementary information and that combined analysis can lead to new insights on on- and off-target effects of small molecules.

## Introduction

Chemo-informatics approaches to rational drug design have traditionally assumed that chemically similar molecules have similar activities. More recently, transcriptional responses of cells treated with small molecules have been used in the lead optimization phase of drug discovery projects^1^ and to reveal similarities among drugs, and quickly transfer indications for drug repositioning.^2-6^

The Connectivity Map (CMAP), the largest peer-reviewed public database of gene expression profiles following treatment of five human cancer cell lines with 1,309 different bioactive small molecules^2, 7^, has been extensively used by both the academic and industrial communities.^3, 8^

Whereas computational medicinal chemistry’s “pros” and “cons” have been extensively addressed over the recent years,^9-17^ on the contrary, the advantages and limits of methods based on transcriptional responses have not been thoroughly addressed.^1, 3^ So far, comparisons of the chemical versus transcriptional “landscape” of small molecules has been performed to elucidate and understanding mode of actions of existing drugs and revealing potential on-label and off-label applications.^18-21^ In this work, on the contrary, we addressed two still unanswered questions: (2) do transcriptional responses and chemical structures provide similar information on the drug mechanism of action and adverse effects? (2) If not, why does the information provided by transcriptional responses and chemical structures differ?

In this work, we compared chemical structures to transcriptional responses in the CMAP dataset by first generating a “structural” drug network by connecting pairs of structurally similar drugs, as measured by 3D pharmacophore descriptors based on Molecular Interaction Fields.^22, 23^ We then compared the structural drug network to a transcriptional drug network where drugs are connected if they induce a similar transcriptional profile.^4, 24, 25^

Through the integrated analysis of chemical structures and transcriptional responses of small molecules, we revealed limitations and pitfalls of both transcriptional and structural approaches, and proposed ways to overcome them. Moreover, we found an unexpected link between drug-induced lysosomotropism and lipid accumulation, common adverse effects, and a specific transcriptional signature mediated by the transcription factor TFEB.

## Results

The CMAP dataset is a collection of transcriptional responses of human cell lines to small molecules. It includes transcriptional profiles following treatment of 1,309 small molecules across five different cell lines, selected to represent a broad range of activities, including both FDA–approved drugs (670 out of 1309 (51%)) and non-drug bioactive ‘‘tool’’ compounds.^2^ An extension of this dataset to more than 5000 small molecules is being completed but it includes only 1,000 genes and it has not been peer-reviewed yet (LINCS http://www.lincscloud.org).^2, 7^ We selected the small molecules present in the CMAP and in the upcoming LINCS resource for a total of 5,452 compounds (**Supplementary Fig. 1**). We then performed a physico-chemical characterization of these 5,452 small molecules by computing 128 physico-chemical descriptors using 3D Molecular Interaction Fields (MIFs) derived from their chemical structures.^26, 27^

Principal Component Analysis (PCA) of the 128 descriptors for all the 5,452 compounds in **Supplementary Figure 2a** reveals that the first two principal components (PC1 and PC2) explain most of the descriptors’ variance (53%). PC1 (36%) is related to descriptors of hydrophobic and aromatic properties (**Supplementary Fig. 2b**), whereas PC2 (17%) to molecular size and shape. Most of these small molecules follow the ‘Rule of Fives (RoFs)’, that is the set of physico-chemical features shared by biologically active drugs: MW ≤500 Da (89%); N.HBA≤10 (93%); N.HBD≤5 (97%); LogP ≤5 (85%) (**Supplementary Fig. 3**).^28,29^

### Chemical structure similarities induce a hierarchical network connecting drugs with similar scaffolds and mode of action

We derived a *structural drug network* where each small molecule is a node and an edge connects two small molecules if they have a similar 3D structures. To this end, we computed the *structural distance* between each pair of small molecules based on the similarity between their 3D-pharmacophore quadruplet-based fingerprints (Methods and **Supplementary Fig. 4**).^30^ A short structural distance (i.e. close to 0) between two compounds indicates that they are structurally similar.

We obtained a symmetric 5,452x5,452 structure-based drug-distance matrix containing 14,859,426 distances between all the possible pairs of drugs. We considered each compound as a node in the network and connected two nodes if their distance was below a threshold value (Methods). The resulting drug network consists of 5,312 nodes and 742,971 edges, corresponding to 5% of a fully connected network with the same number of nodes (14,859,426 edges) (http://chemantra.tigem.it). We subdivided the network into communities consisting of groups of densely interconnected nodes by means of the Affinity Propagation (AP) clustering algorithm ^31, 32^ on the network matrix (Methods).^4^ We identified 288 communities (containing more than 3 drugs) across 5,302 drugs (out of 5,452) that group together compounds sharing similar chemical functionalities, scaffolds and sub-structural fragments. The AP clustering assigns to each community an “exemplar”, i.e. the drug whose structure best represents the structures of the other drugs in the community. By iteratively applying the AP clustering on the exemplars, we could further group communities into 42 *Rich Clubs, i.e. clusters* of drug communities that are structurally related but with distinct characteristic functional groups (**Fig. 1**).

**Figure 1:**
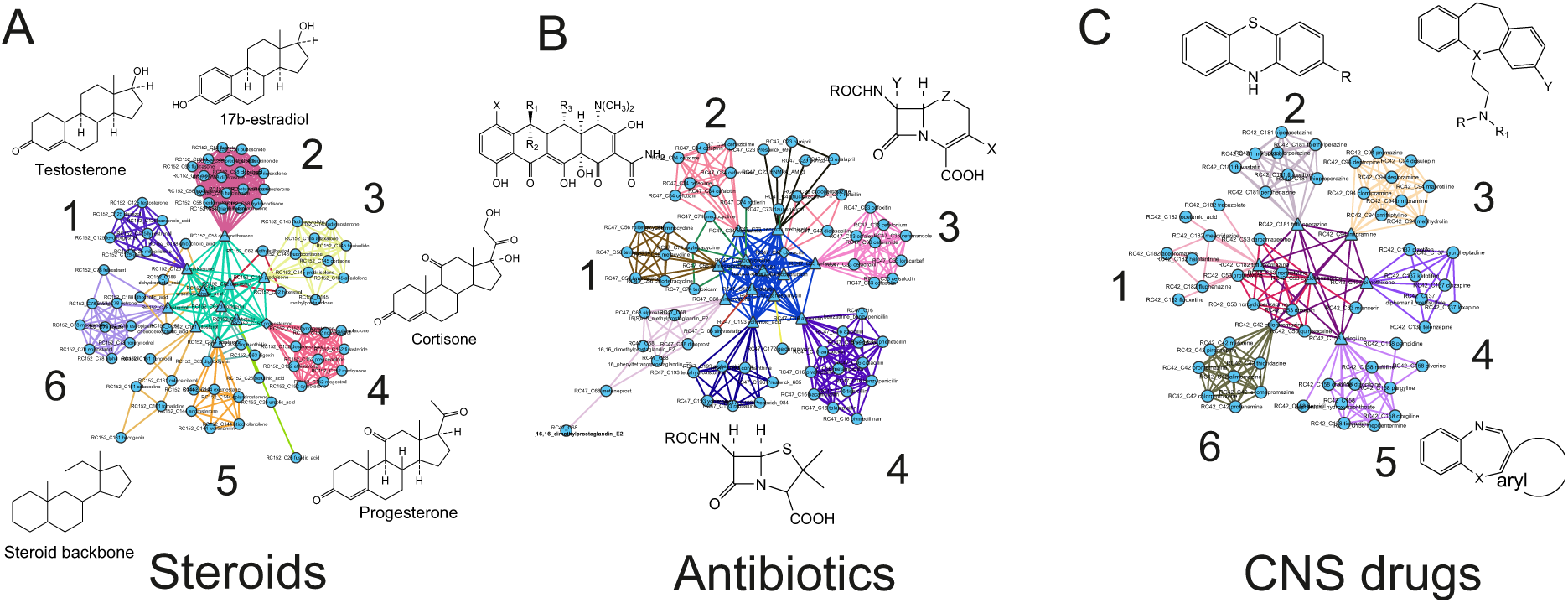
The structural network among 5452 compounds. The network is partitioned into *communities* (groups of highly interconnected nodes) and *rich-clubs* (groups of communities) sharing common chemical structures and enriched for drugs with similar Mode of Action. Examples of three Rich Clubs are shown: **a**) The steroids rich-club (1: testosterone scaffold, 2: estradiol scaffold, 3: cortisone scaffold, 4: progesterone scaffold, 5 and 6: mixed steroids); **b**) The antibiotics rich-club (1 and 2: tetracycline scaffold, 3: cephalosporin scaffold, 4: penicillin scaffold); **c**) The CNS-acting drug rich-club (1 and 2: phenothiazine scaffold, 3-6: various tricyclic antidepressant scaffolds).

To assess the structural network, we collected the ATC (Anatomical Therapeutic Chemical) code, an alphanumerical hierarchical pharmacological classification, for 936 out of 5452 drugs (Methods). We then verified that drugs connected in the network tend to share the same ATC code (**Supplementary Fig. 5**). We also verified that drugs within a community share a common therapeutic application. Indeed, 230 out of 288 (80%) structural communities were significantly enriched for compounds sharing the same ATC code (p-values <0.05) (**Supplementary Fig. 6**). These results demonstrate that inspection of the structural drug network can provide useful information on the drug mechanism of action and possibly help in identifying candidates for drug repositioning.

### Chemical similarity between drugs is largely uncorrelated with similarity in induced transcriptional responses in CMAP

In a previous study^4^,^24^ we reported on the construction of a “transcriptional network” among 1,309 small-molecules part of the CMAP dataset^2^ (http://mantra.tigem.it) where two drugs are connected by an edge if they induce a similar transcriptional response. Briefly, in CMAP each transcriptional response is represented as a list of genes ranked according to their differential expression in the drug treatment versus control. Since each drug is associated to more than one ranked list (cell, dosage, etc.), to obtain the transcriptional network, we first computed a Prototype Ranked List (PRL) by merging together all the ranked lists referring to the same compound following the Borda Merging method to generate a single ranked list^4^. The PRL thus captures the consensus transcriptional response of a compound across different experimental settings, consistently reducing non-relevant effects due to toxicity, dosage, and cell line.^4^ Transcriptional similarity was then quantified among the 1,309 PRLs (one for each drug) by Gene Set Enrichment Analysis and represented as a distance (i.e. 0 for identical responses, and greater than 0 if dissimilar)^4^. The transcriptional network was obtained by connecting two nodes if their distance was below a significant threshold value chosen so that the total number of edges is equal to 5% of a fully connected network with the same number of nodes (856,086 edges).

Here, we compared structural and transcriptional similarities among all pairs of drugs, part of the CMAP dataset, as shown in **Figure 2**, where each point is a drug-pair and its position in the plane represents the structural (x-axis) and transcriptional (y-axis) distance between the two drugs, for a total of 856,806 drug-pairs. The structural-transcriptional plane can be subdivided into four quadrants by straight lines representing the significance thresholds for the transcriptional (y-axis) and structural (x-axis) distances: quadrant I (5.1% of drug-pairs) contains drug-pairs with similar structures but inducing different transcriptional responses; quadrant II (0.3% of drug-pairs) contains coherent drug-pairs that are both structurally and transcriptionally similar; quadrant III (4.0% of drug-pairs) consists of drug-pairs with different structures but inducing similar transcriptional responses; finally drug-pairs different both in structure and transcription are found in quadrant IV (91% of drug pairs). This quadrant contains most drug-pairs since two random drugs usually have no common function at all. We call drug-pairs in quadrant I and III *incoherent* because of the discrepancies between structural and transcriptional similarities, whereas drug-pair in quadrant II and IV are *coherent*.

**Figure 2:**
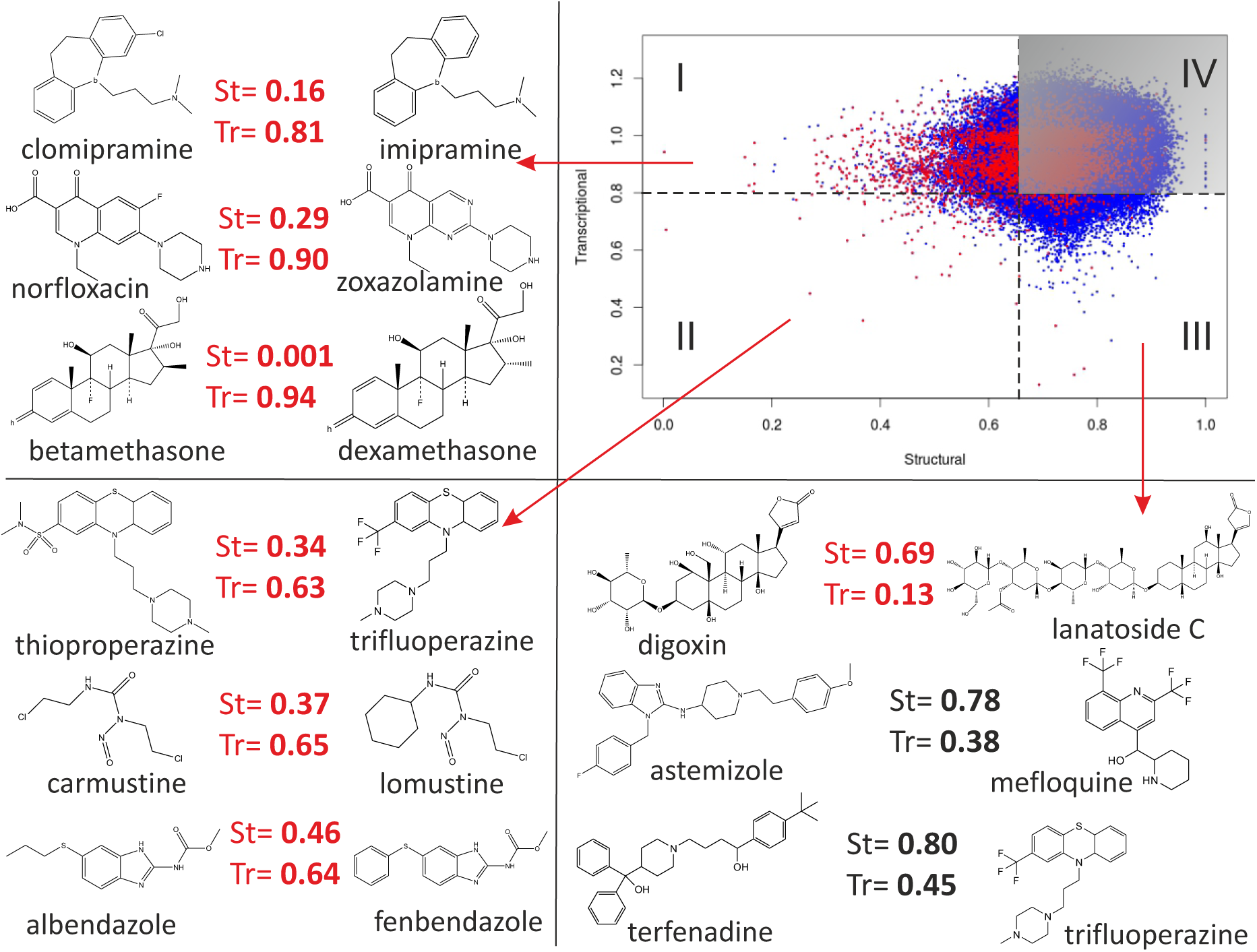
Comparison of transcriptional and structural distances between 784 CMAP compounds having at least one ATC annotation. Each dot represents the structural (x-axis) and transcriptional (y-axis) distance between two compounds. A total of 306,936 drug-pairs are shown. <u>Drug-pairs having the same clinical application as annotated by their ATC code are represented by red dots</u>. Dashed lines represent the significance threshold for the transcriptional (horizontal line) and structural (vertical line) distance, splitting the plane into four quadrants. Representative examples of drug-pairs are shown for quadrants I, II and III: drug-pairs in quadrant I have similar structure but induce different transcriptional responses; drug-pairs in quadrant II exhibit both similar structure and similar transcriptional responses; drug-pairs in quadrant III have different structures but induce similar transcriptional responses.

Overall, **Figure 2** shows that the information detected by transcriptional responses and chemical structures tend to be different and independent of each other. We therefore decided to investigate the causes for this lack of correlation.

### Chemically similar drugs do not induce similar transcriptional responses because of weak transcriptional effects

Drug pairs sharing highly similar chemical structures but very different transcriptional responses are found in **Figure 2** (quadrant I). These drug-pairs exhibit an unexpected behaviour, since they are chemically similar molecules with the same therapeutic application (i.e. ATC code) but inducing very different transcriptional responses.

The most surprising example was the betamethasone/dexamethasone drug-pair in **Figure 2** (quadrant I). Both drugs are glucocorticosteroids binding the Glucocorticoid Receptor (GR) with very high affinity and nearly identical in structural since they are enantiomers of each other. Transcriptionally, on the contrary, these two drugs appear to be completely different (**Supplementary Fig. 8g**).

One possible explanation is that these compounds cause weak transcriptional effects in the cell lines used in CMAP, and thus the measured transcriptional responses are too noisy to be informative.

To assess whether a perturbation (e.g. drug treatment) leads to a strong and informative transcriptional response, we introduce the “transcriptional variability” score (TV). The TV score is based on the assumption that when the cellular context contains the necessary molecular *milieu* to make it responsive to a small molecule, then multiple treatments with the same compound will yield consistent and similar transcriptional responses.

We computed the TV of a small-molecule as follows: given M transcriptional responses to the same small-molecule in the same cell line (i.e. ranked list of differentially expressed genes as in CMAP), we evaluate the transcriptional distances between all the M(M-1)/2 pairs of transcriptional responses and then take their median value as a measure of TV (if M = 2 then TV is defined as the maximum distance). A TV close to 0 implies very similar transcriptional responses across replicates, indicating that the small molecule induces a reliable transcriptional response across all the experiments. On the contrary, a high TV implies very different transcriptional responses across replicates, hence a weak and unreliable transcriptional signature.

To assess whether TV is indeed able to detect informative versus non-informative transcriptional responses to small-molecules, we exhaustively computed the TV of 1165 CMAP drugs (out of 1309) for which at least two transcriptional responses in the same cell line were available (**Supplementary Table 1**). Out of the 1165, 858 (73%) have a TV score greater than the significance threshold implying that most drugs in CMAP induce a weak transcriptional response (Methods).

We compared the TV of drugs belonging to different classes, which were chosen because of their expected activity, or lack thereof, in the CMAP human cancer cell lines (**Fig. 3 and Supplementary Table 1**). As expected, glucocorticosteroids exhibit higher values of TV when compared to the other classes of drugs. Similarly, antibiotics and NSAIDs induce very weak transcriptional responses (high TV values). Indeed, antibiotics target bacteria-specific proteins, whereas NSAIDs act on cell-specific enzymatic pathways with marginal effects on transcription. Most antihistamines and antipsychotics induce weak transcriptional responses since they target specific cell membrane receptors lowly, or not expressed, in CMAP cancer cell lines and with no direct transcriptional effects.

We observed that drugs with a high TV, hence exhibiting a weaker transcriptional response, tend to have higher transcriptional distances from the other drugs in CMAP (i.e. they tend to be isolated in the network) and vice-versa (**Supplementary Fig. 7**). Consistently with this observation, compounds within these drug-classes tend to be found in drug-pairs belonging mostly in quadrant I (structurally similar and transcriptionally different) and quadrant IV (structurally and transcriptionally different) as shown in **Supplementary Fig. 8**.

**Figure 3:**
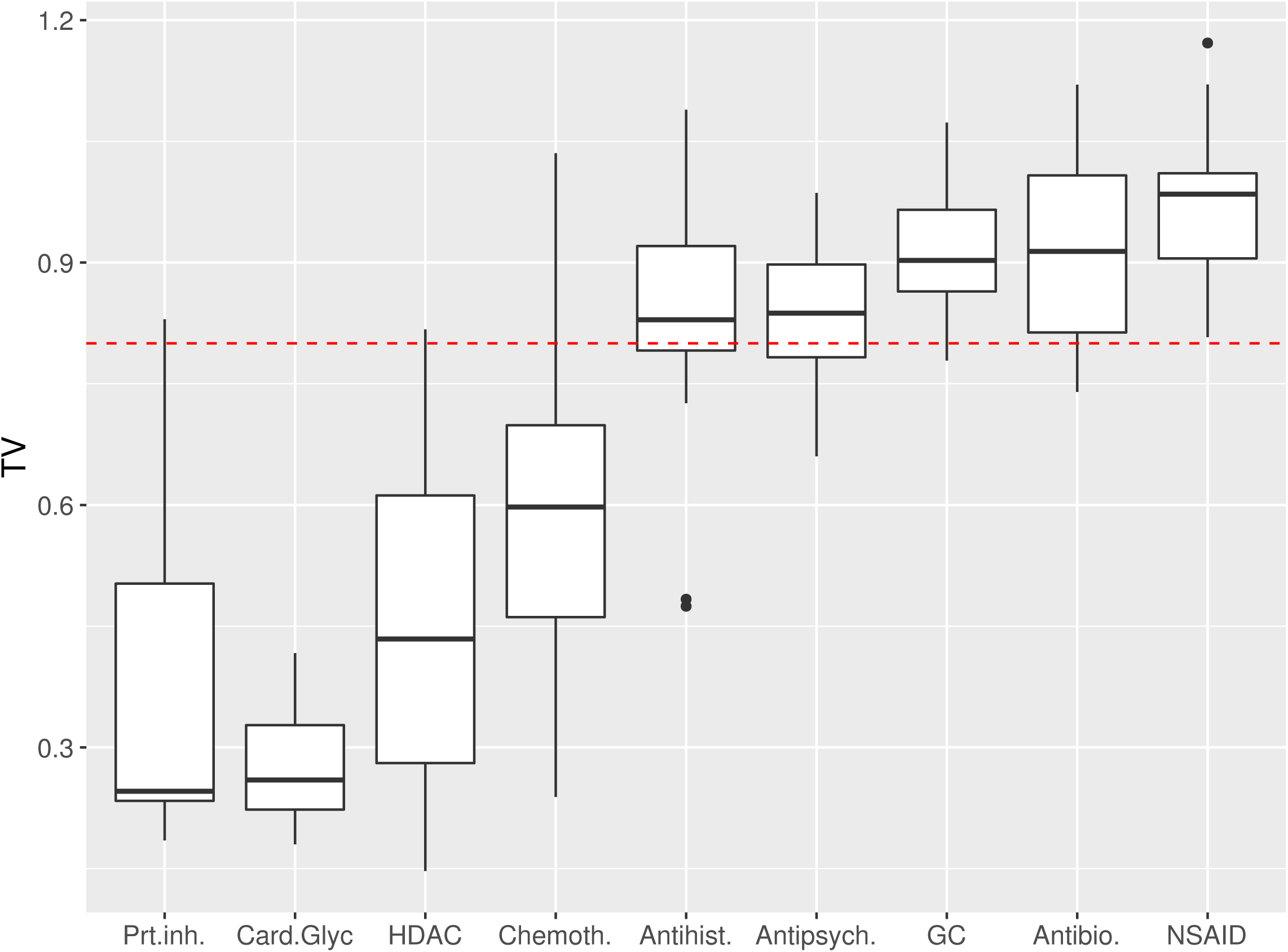
The Transcriptional Variability (TV) of different drug classes. Box-plots summarising the TV for drugs within each class. The bold line in each box represents the median, while the whiskers represent the 25^th^ and the 75^th^ percentile. Dots represent outliers. Prt.inh.: Protein synthesis inhibitors; HDAC: histone deacetylase inhibitors; Chemoth.: chemotherapeutic agents; Antibio.: antibiotics; NSAIDs: non-steroid antinflammatory agents; GC: glucocorticoids; Antipsych: antipsychotics; Antihist: antihistamines.

Conversely, drugs with the lowest TV (**Fig. 3 and Supplementary Table 1**), and thus with strong transcriptional responses, consist mostly of lipophilic molecules acting as protein synthesis inhibitors, chemotherapeutic drugs and other DNA/RNA intercalating agents, and histone deacetylase inhibitors, which all have a strong activity in most cell types (**Supplementary Fig. 9**). Interestingly, several cardiac glycosides were also found to have a low TV. As shown in **Supplementary Fig. 8**, in this case drug-pairs consisting of compounds in these drug-classes tend to be found in quadrant III (structurally different but transcriptionally similar).

### Removing weak transcriptional responses from the CMAP dataset improves drug classification performances

We reasoned that by removing drugs with a high TV, the performance of computational approaches based on gene expression to elucidate the MoA of a drug should improve.^4, 24^ We thus partitioned the small molecules included in CMAP in two sets according to their TV score, obtaining a high-TV set and a low-TV set with the same number of drugs to facilitate the comparison. We then assessed the performance of the transcriptional distance between two drugs in correctly identifying those pairs sharing the same therapeutic application (i.e. the same ATC code), when using either drugs in the high-TV set or those in the low-TV set, as previously described.^4^ As shown in **Figure 4**, the low-TV set performance far exceeds the high-TV set performance, which is almost random. Moreover, the correlation between structural distance and transcriptional distance in the chemical-transcriptional landscape of small molecules in **Figure 4** increases if only drugs in the low-TV set are used (**Supplementary Fig. 10**).

**Figure 4:**
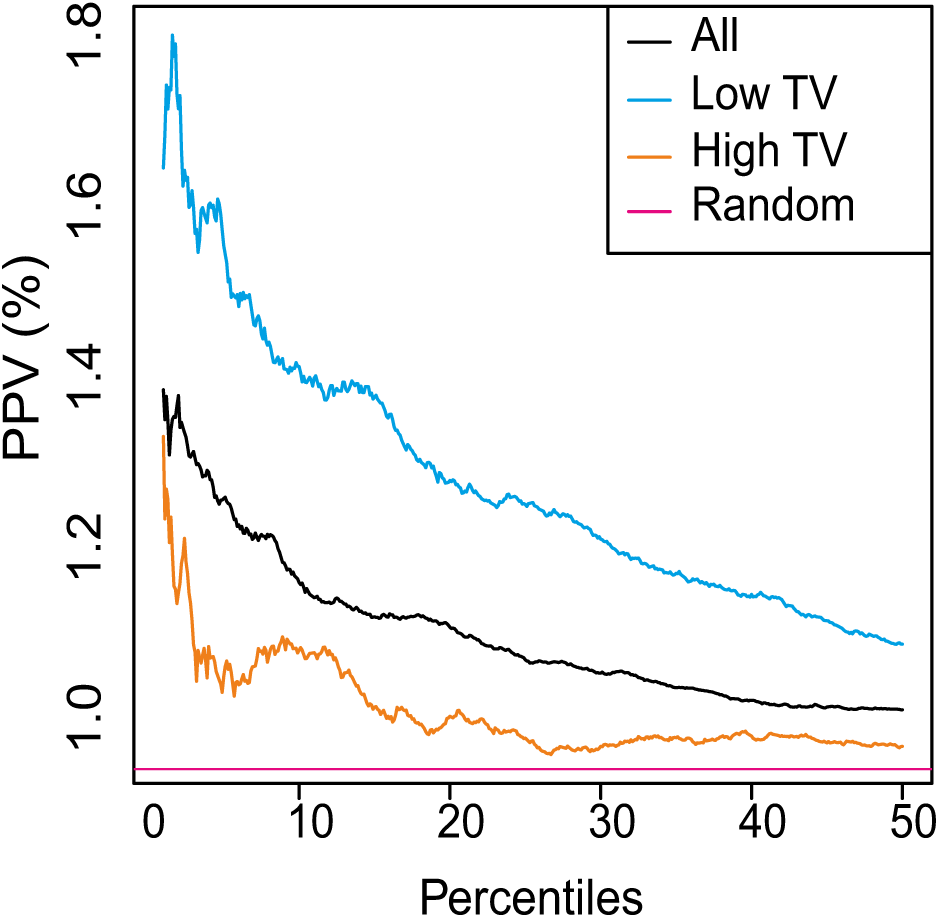
Performance of the transcriptional distance in detecting drugs with the same ATC code. Compounds were divided into three sets: (All) the 1165 compounds in CMAP having at TV value; (High TV) 582 compounds with a TV higher than the median TV among all the compounds; (Low TV) 582 compounds with a TV lower than the median TV. For each set, the transcriptional distance of each drug-pair was computed. Drug-pairs were then sorted according to their transcriptional distance, with drug-pairs with the smallest distance towards the origin of the *x-axis;* the Positive Predictive Value (PPV) was computed as the percentage of True Positives over False Positives plus True Positives and shown on the *y-axis*. The PPV obtained by randomly sorting drugs is also shown (Random).

Overall, these results show that the TV score can discriminate between informative and non-informative transcriptional responses that result from the activity, or lack thereof, of small molecules in a specific cell line.

### Drugs with different chemical structures and modes of action may induce similar transcriptional responses related to lysosomal stress and phospholipidosis

**Figure 2** (quadrant III) includes drug-pairs with very different molecular structures but which are transcriptionally similar. We identified two obvious causes for the discrepancy between transcriptional and structural similarities: (i) most of the drug-pairs in this region have at least one drug with a very large size (usually a natural compound) (**Fig. 2 and Supplementary Fig. 11**) hence, global chemical similarity metrics, such as the one used here, may fail; (ii) the direct molecular targets of two drugs in a pair may be different but act in the same pathway (e.g. purine synthesis inhibitors methotrexate/mycophenolic-acid that act on different molecular targets but both block DNA synthesis, **Supplementary Fig. 11**) ^33-35^.

Figure 2 (quadrant III), however, contains also a large fraction of drug-pairs that are not large molecules and do not act in the same pathway, nor share the same therapeutic application, but nevertheless have very similar transcriptional profiles. To investigate why this is the case, we ranked drug-pairs in this quadrant by their transcriptional distance in ascending order (**Supplementary Table 2**). We noticed that the top-ranked most transcriptionally similar drug-pairs included well known “lysosomotropic agents” inducing large vacuolization in cells such as astemizole, terfenadine and mefloquine (**Table 1**).^36-38^ Among these agents, astemizole and terfenadine are no longer in use because of cardio-toxicity caused by their potassium channel blocker activity (hERG encoded by *KCNH2*), which may lead to fatal cardiac arrhythmia.^39, 40^ The lysosomotropic effect of these small molecules has been attributed to their ability to cross lysosomal membrane and remain trapped within the lysosome by a mechanism known as pH partitioning.^41 42-44^ Most lysosomotropic agents belong to the class of cationic amphiphilic drugs (CADs) containing both a hydrophobic and a hydrophilic domain. CADs have increased probability to cause drug-induced phospolipidosis (PLD),^45^ a lysosomal storage disorder characterized by the accumulation of phospholipids within the lysosome by unclear molecular mechanisms, leading to cellular stress.^46-50^ Indeed among the lysosomotropic drugs involved in the most transcriptionally similar drug-pairs (**Table 1**), there were also three known PLD inducing drugs (astemizole, suloctidil and trifluoperazine).

**Table 1:** Drug-pairs with different chemical structures but inducing very similar transcriptional responses. Drug-pairs in **Figure 2** (quadrant III) were ranked by transcriptional distance (Tr. Dist.). Only the top 20 ranked drugs pairs are shown together with their structural distance (Str. Dist.). Lysosomotropic drugs are shown in italic and phospholipidosis inducing drugs in bold.

| Drug A | Drug B | Tr. Dist. | Str. Dist. |
| --- | --- | --- | --- |
| digoxin | lanatoside_C | 0.131 | 0.693 |
| digoxin | proscillaridin | 0.166 | 0.758 |
| lanatoside_C | proscillaridin | 0.187 | 0.776 |
| rifabutin | vorinostat | 0.286 | 0.826 |
| **astemizole** | *terfenadine* | 0.337 | 0.724 |
| **astemizole** | *mefloquine* | 0.385 | 0.776 |
| doxorubicin | mitoxantrone | 0.414 | 0.651 |
| *mefloquine* | *terfenadine* | 0.421 | 0.767 |
| chlorzoxazone | clindamycin | 0.442 | 0.829 |
| chlorzoxazone | glibenclamide | 0.445 | 0.791 |
| *terfenadine* | **trifluoperazine** | 0.453 | 0.758 |
| irinotecan | *phenoxybenzamine* | 0.455 | 0.800 |
| **suloctidil** | *terfenadine* | 0.466 | 0.696 |
| astemizole | **trifluoperazine** | 0.469 | 0.718 |
| *protriptyline* | **trifluoperazine** | 0.472 | 0.713 |
| niclosamide | **trifluoperazine** | 0.472 | 0.688 |
| *mefloquine* | **trifluoperazine** | 0.478 | 0.674 |
| doxazosin | sulconazole | 0.481 | 0.776 |
| lomustine | *phenoxybenzamine* | 0.484 | 0.719 |

We hypothesised that “lysosomotropic” stress induced by these compounds could explain their similarity in transcriptional responses. We therefore selected 187 CAD compounds present in CMAP according to their physico-chemical properties (LogP >3; pKa >7.4)^43^. Within these CAD compounds, we searched the literature for lysosomotropic drugs known to induce PLD,^45^ which, according to our hypothesis, should elicit a strong transcriptional response. We thus identified a total of 35 compounds (PLD/CAD) (**Supplementary Table 3**).

We verified that PLD/CAD compounds tend to induce a stronger transcriptional response (i.e. a lower TV) (**Supplementary Fig. 12**) and they tend to be transcriptionally similar among them (but not structurally) despite having different mode of action and therapeutic applications (**Supplementary Fig. 13**).

We next asked which genes were transcriptionally modulated by the majority of PLD/CAD compounds. We performed Drug Set Enrichment Analysis (DSEA),^51^ a computational approach we recently developed to identify gene-sets that are transcriptionally modulated by most drugs in a given set. The most significant gene-set shared by the 35 PLD/CAD compounds, out of about 5,000 gene-sets within the Gene Ontology (GO) database, was the GO-Cell Component term “lysosome” consisting mainly of genes coding for lysosomal enzymes and ion channels (p=5.03x10^-8^ – **Supplementary Table 4**), thus in agreement with the “lysosomotropic” effect of these drugs.

Recently, the transcription factor E-box (TFEB) has been found to be a major player in the transcriptional control of lysosomal genes in response to a variety of cellular and environmental stresses.^52^ In normal nutrient conditions TFEB is phosphorylated by the mTORC1 complex on the lysosomal surface. This phosphorylation favours TFEB binding to 14-3-3 proteins and its retention in the cytoplasm.^53-55^ Upon stress signal, such as nutrient deprivation, mTOR is inhibited, the calcium-dependent phosphatase Calcineurin is activated, and TFEB is de-phosphorylated shuttling to the nucleus where it transcriptionally controls lysosomal biogenesis, exocytosis and autophagy.^53-59^ Moreover, TFEB was shown to translocate to the nucleus upon amiodarone treatment, a well known lysosomotropic agent.^60^ We thus decided to investigate whether TFEB activation was responsible for the characteristic transcriptional response induced by PLD/CAD compounds.

### The transcriptional response of PLD-inducing compounds is mediated by TFEB

We performed a panel of High Content Screening assays including the TFEB nuclear translocation assay (TFEB-NT)^59^ at 3h and 24h following drug administration at different concentrations (0.1μM, 1μM and 10μM) for 34 out of 35 PLD drugs (1 drug was not available to us at the time). Additional HCS assays at 24h included LAMP-1 immunostaining and Lysotracker dye to quantify lysosomal compartment (Methods), GM130 and PDI immunostaining to detect morphological changes in the Golgi and ER (Endoplasmic Reticulum) compartments, both of which have been recently suggested to be involved in PLD aetiology (Methods). We also performed the LipidTox assay at 48h to check for the accumulation of phospholipids to confirm PLD at least in vitro (Methods).

Quantification of the HCS assays for the 34 PLD drugs are reported in **Supplementary Figure 13** and Supplementary **Table 5**. Nuclear translocation of TFEB at 3h was observed for 18 out of 34 drugs (53%) increasing to 29 drugs at 24h (85%). Out of these 29 drugs, 27 induced an increase in lysosome size and number as evidenced by LAMP1 and Lysotracker staining, and all 29 drugs induced accumulation of phospholipids according to the Lipidtox assay (100%). Only 5 drugs did not induce TFEB translocation at 24h, and just 1 out of these 5 drugs was positive in the Lysotracker assay, while 4 of them were positive in the Lipidtox assay. None of the drugs tested were positive for the Golgi marker and only 6 were positive for the ER marker, albeit marginally.

Overall, HCS confirmed a concentration dependent nuclear translocation of TFEB for 29 out of 34 drugs (85% at 24h) with a concomitant perturbation of the lysosomal compartment for 28 out of 34 drugs (82%) occurring mostly at the highest dosage tested (10 μM). Furthermore, HCS revealed an accumulation of lipid in vitro at 48h following treatment with the 34 drugs (100%) at the highest dosage tested (10 μM), as previously reported in the literature.^45^

These results support the role of TFEB in shaping the transcriptional response of cells treated with PLD inducing drugs in a way completely unrelated to their MoA. We next asked whether the activation of TFEB (or TFE3, another member of the MiT family of transcription factors with similar functions) is a consequence of lysosomal stress upon compound treatment or if it is directly related to the induction of the PLD phenotype. Thus, we set up a HCS Lipidtox assay using TFEB wt versus TFEB/TFE3 KO in HeLa cell type, administering high dosage of chloroquine (50 μM) known to induce lipids accumulation in cells at 48h. **Supplementary Fig. 15a, b** show no major differences in terms of spot intensity in the Lipidtox assay, thus confirming that TFEB activation is a consequence of lysosomal stress and not an inducer of PLD.

As this manuscript was under review, Lu et al reported an increase in TFEB, TFE3 and MITF translocation to the nucleus in ARPE-19 cells together with lysosomal activation and lipid accumulation following treatment with 8 lysosomotropic compounds, well in agreement with our results. ^50^

### A PLD-specific transcriptional signature can predict compounds inducing lipid accumulation

We combined the transcriptional responses elicited by the 35 PLD/CAD compounds into a consensus transcriptional response (“PLD” signature) and computed its transcriptional distance from all the other 1274 (i.e. 1309-35) CMAP compounds (Methods). We reasoned that drugs inducing a transcriptional profile similar to the PLD signature should have a higher probability of inducing lipid accumulation than the other drugs. Surprisingly, 258 compounds out of 1274 (20%) cMAP compounds were found to be similar to the PLD signature (**Supplementary Table 6**). About a third of these drugs are CADs (77 out of 258 (30%)).

**Figure 5** reports a breakdown by ATC classes of drugs for which an ATC code was available and that were found to induce a transcriptional response similar to the PLD signature. Some drug classes (ATC classes N05, N06 and R06 including antihistamines and antipsychotics) are enriched for known PLDs^45, 47^. Other classes cause global cellular stress responses not mediated by their physico-chemical properties, but rather because of their direct molecular targets, such as anticancer compounds that block cell-cycle (e.g. ATC class L01 composed of CDK2 and Topoisomerase I, II inhibitors). Antihelmintics (ATC P02) and antifungals (ATC D01), despite being neither CADs nor PLDs, were also found among the PLD node’s neighbours. Several recent reports in the literature have found antihelmintics to induce an anti-proliferative effect in cancer cell lines by indirectly inhibiting the mTOR pathway thus inducing TFEB activity, which may explain their PLD-like transcriptional response.^55, 60-63^ Calcium channel blockers were also found to induce a transcriptional response similar to PLDs, which may be expected since calcium signalling has been involved in autophagy regulation and lysosomal function.^59^ Interestingly, some cardenolides (ATC C01 and C07) were also found to contain the PLD signature, despite not being CADs (median distance equal to 0.71).^64, 65^

**Figure 5:**
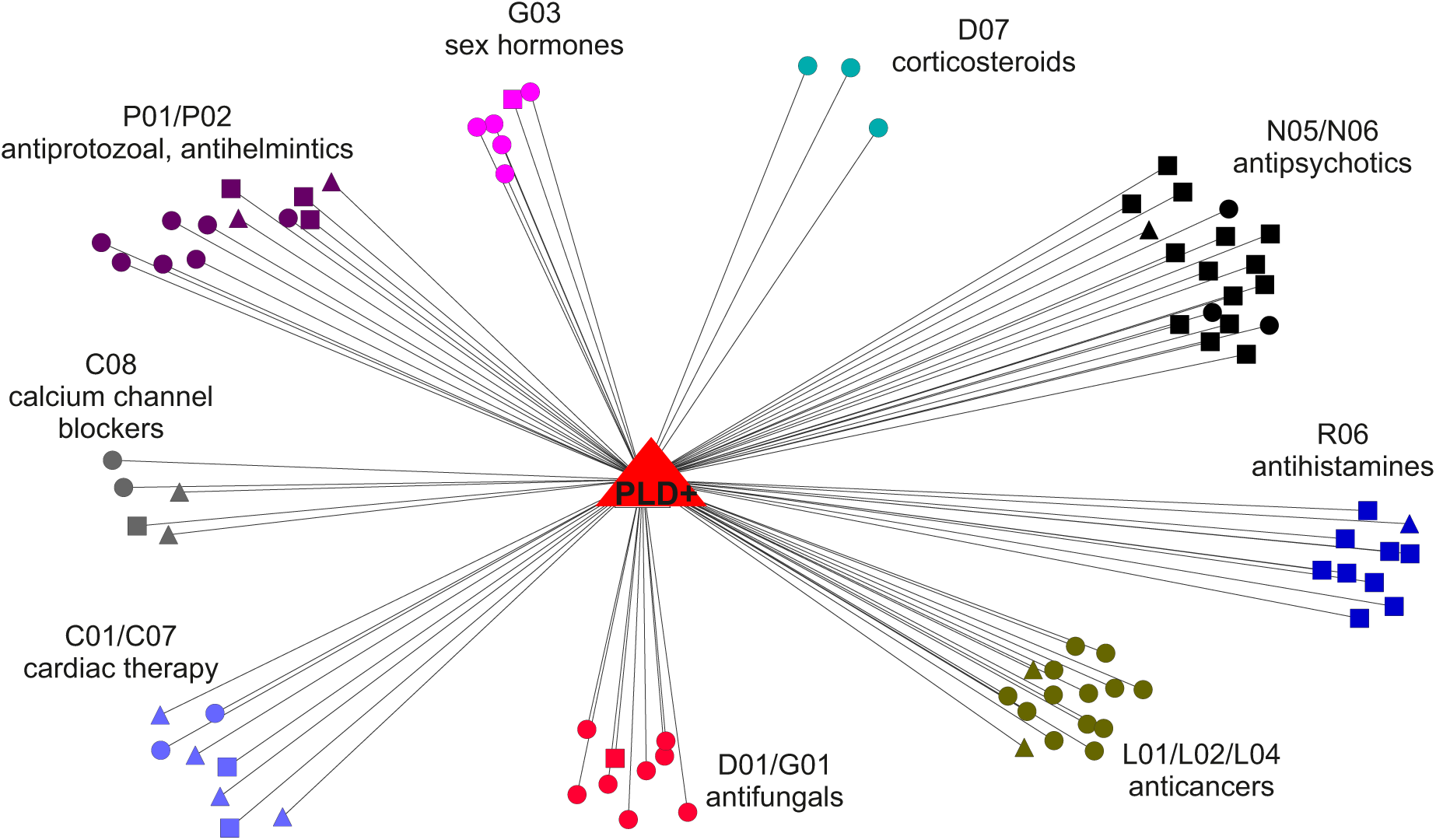
Drugs inducing a lysosomotropic gene expression signature. The transcriptional responses elicited by 8 lysosomotropic compounds were combined into a single node in the transcriptional drug network (red triangle). Transcriptional distances to this lysosomotropic gene expression signature were computed for all the 1309 drugs in CMAP. Only drugs with a transcriptional distance below the significance threshold are shown (0.8) and colour-coded according to their ATC classification.

To experimentally validate the usefulness of the PLD transcriptional signature in identifying novel PLD drugs, we selected the top quartile of the 258 drugs (i.e. 25% of 258=64 drugs) with the shortest transcriptional distance to the PLD node and performed HCS for lipid accumulation following drug treatment at three different concentrations (Lipidtox assay) (**Supplementary Table 7**). Twenty-two out of the top 64 small molecules were present in our HCS small-molecule library. Overall 11 out of 22 (50%) compounds were positive to the Lipidtox assay (**Supplementary Table 7**), including Terfenadine, a cardiotoxic lysosomotropic CAD, not reported to be a PLD inducer in the literature, which caused a strong accumulation of lipids, as shown in **Figure 6** (LipiTox Intensity Spot: 450.93 at a concentration of 10μM).

**Figure 6:**
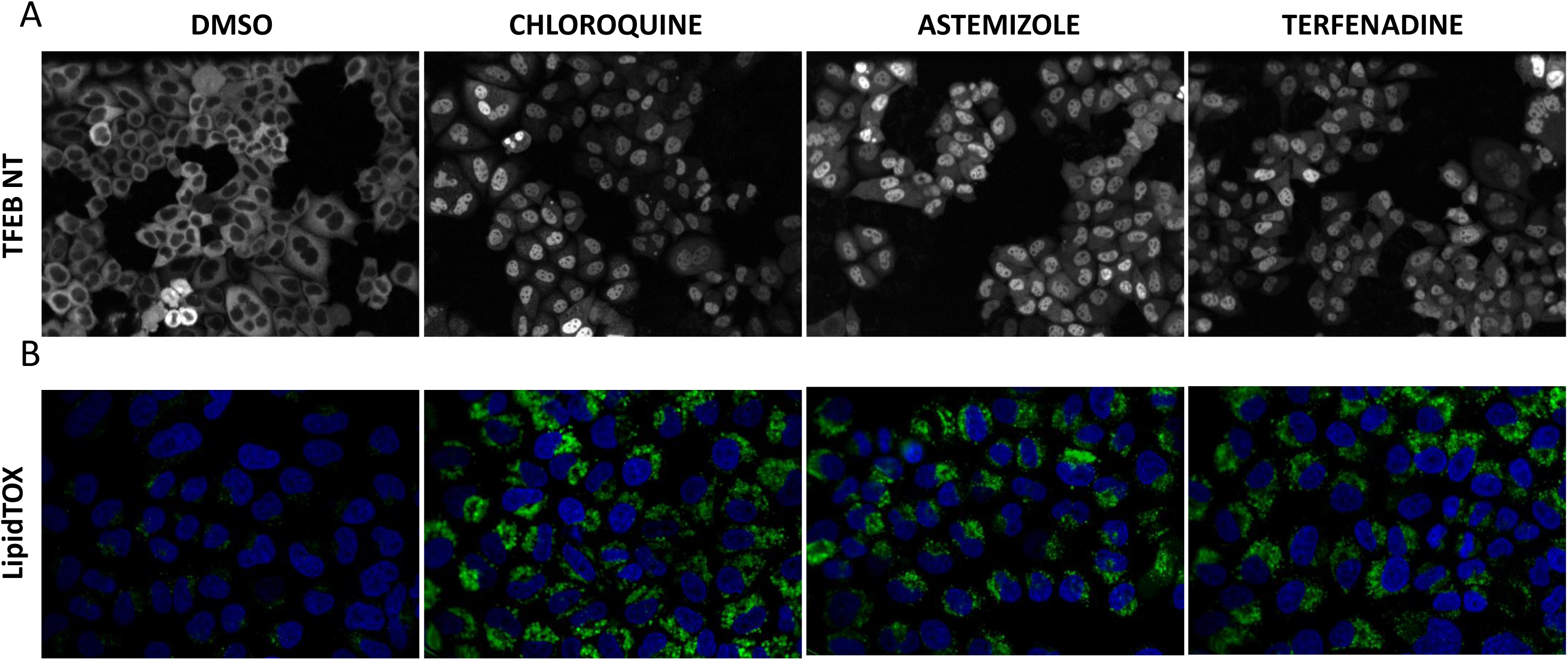
Effects of drugs on TFEB nuclear translocation and LipidTOX assay. A) TFEB localization in stably HeLa cells overexpressing TFEB-GFP and treated with DMSO or the indicated drugs. B) Lipid accumulation in HeLa cells was detected by staining with LipidTOX reagent upon drug treatment.

Overall, our data demonstrate the value of the PLD transcriptional signature in identifying compounds potentially inducing lysosomal stress and phospholipidosis.

### The PLD transcriptional signature affects transcriptional responses to drug treatment in a concentration dependent manner

We next investigated whether the PLD expression signature was linked to the elevated drug concentration used in the CMAP experiments, in agreement with the HCS results indicating a dose dependent TFEB nuclear translocation (**Supplementary Fig. 14**). Indeed 5,747 out 6,100 CMAP gene expression profiles (94%) were measured at high drug concentrations ranging from 1μM to 10 mM, while the remaining 353 (6%) at lower concentrations ranging from 10nM to 0.5 μM. We thus searched CMAP for PLD-inducing drugs for which both high and low concentration instances were present. We selected 5 drugs (out of 35 PLD) drugs: raloxifene (ER antagonist at 0.1 μM and 7.8 μM), tamoxifen (ER antagonist at 1 μM and 7.0 μM), amitriptyline (antidepressant 1 μM and 12.8 μM), thioridazine (antipsychotic at 1 μM and 10 μM) and chlorpromazine (antipsychotic at 1 μM and 11.2 μM). We then generated two additional transcriptional responses (LOW and HIGH) for each of these 5 drugs by analysing separately the low and high concentration experiments (**Methods, Supplementary Figure 15** and **Supplementary Table 8**).

The HIGH transcriptional responses for the 5 drugs were more similar to the PLD signature than the corresponding LOW transcriptional responses (**Supplementary Table 8**), confirming an increased alteration of the transcriptional response caused by high drug dosages. Moreover, the HIGH transcriptional responses of 4 out of 5 drugs were connected to a much larger number of drugs in the transcriptional network when compared to their LOW transcriptional response counterparts (**Supplementary Fig. 16**). Raloxifen, a selective estrogen receptor modulator (SERM), is the only drug tested also at sub-micromolar concentrations (0.1 μM). When using the HIGH transcriptional response, raloxifene is predicted to be transcriptionally similar to 154 compounds (**Supplementary Fig. 16 and Supplementary Table 8**), none of which behaving as a SERM, with the most similar being trifluoperazine, an antipsychotic drug with known PLD-inducing properties. On the contrary, when the LOW transcriptional response is used, raloxifene is predicted to be transcriptionally similar only to 4 compounds, the most similar one being tamoxifen, a well-known SERM.

## Discussion

By analysing a large set of chemical structures, we generated a network representing structural similarities among compounds that can be used to automatically group together drugs with similar scaffolds and mode-of-action. Other methods to cluster drugs based on structural similarity have been proposed in the literature^16^ but no hierarchical classification of drugs in communities and rich-clubs based on the network structure has been previously performed. By comparing the structural drug network with the transcriptional drug network, we observed broad differences between the two: drugs can be very similar in terms of the transcriptional response they induce, but with unrelated chemical structures, or vice-versa have very similar structures but induce diverse transcriptional responses.

Here, we identified a set of confounding factors that can hinder the usefulness of transcriptional based methods. We introduced a simple but powerful measure, “Transcriptional Variability” (TV), to assess the strength and robustness of the transcriptional response of a cell to a drug treatment.

In the original CMAP study,^2^ the authors indeed recognised that although gene-expression signatures can be highly sensitive, they may be uninformative if measured in cells that lack the appropriate physiological or molecular context, but offered no solution to identify such cases. We observed that glucocorticoids tend to have a high TV, hence uninformative transcriptional profiles. Indeed, MCF7,^66, 67^ HL60 and PC3^68, 69^ cell lines used in CMAP may exhibit resistance to glucocorticosteroids^2^. Hence, if not filtered out, computational analysis of their transcriptional responses may be misleading and lead to wrong conclusions, e.g. such that betamethasone and dexamethansone have a different mode of action (**Figure 2**).

We also uncovered a transcriptional signature common to a subset of transcriptionally similar but structurally distinct drugs profiled in CMAP that is not related to their mode of action, but rather to cellular toxicity caused by lysosomal stress and lipid accumulation.

We further demonstrated by HCS that PLD inducing drugs have little effect on ER and Golgi morphology, but rather increase the number and size of lysosomes, as previously reported in the literature, and induce the nuclear translocation of the transcription factor TFEB, a master regulator of lysosomal biogenesis and autophagy. We show that the transcriptional signature present in the transcriptional response of PLD inducing drugs is mainly driven by TFEB activation. These results may help in further elucidating the effect of lysosomotropic PLD-inducing drugs on autophagy.^70^ Moreover, the PLD transcriptional signature may be a useful tool for identifying and repositioning drugs as inducers of TFEB activation and thus of authophagy.^57^

Our findings are relevant for all those studies relying on CMAP transcriptional responses to determine drug mode of action and for drug repositioning. Here, we show that very high and not physiological compound concentrations, such as the ones used in the CMAP dataset, increase the chance of off-target effects including lysosomotropism and phospholipidosis. Somewhat surprisingly, despite the high concentrations used, only a minority of compounds in CMAP (~30%) have reproducible transcriptional responses (TV<0.8). Notwithstanding these limitations, the CMAP still contains relevant information on drug activity if properly analysed, allowing to correctly discriminate among different classes of drugs^3^ and it can provide complementary information to that obtained by HCS.^4, 71-73^

Based on the results here presented, we suggest guidelines to prevent inconsistencies and erroneous conclusion when using transcriptional responses of small molecules for drug discovery and drug repositioning: (i) the transcriptional response elicited by a drug can be uninformative. Hence these responses must be detected and then excluded from further analyses. We demonstrated that this can be achieved by assessing the Transcriptional Variability (TV) of the drug–induced transcriptional response across multiple replicates; (ii) drug treatment can cause cellular stress unrelated to the drug MoA and thus affect the drug-induced transcriptional response by partially masking transcriptional changes directly related to the drug molecular targets. We generated a PLD transcriptional signature which can be used to detect these compounds. This signature is particularly strong if drug concentrations used to treat cells are above their clinically relevant concentrations. One way to avoid this is to use clinically relevant (sub-micromolar) concentrations; (iii) in the case of natural compounds, computational approaches based on transcriptional responses maybe more informative than those based on structural approaches, because of the large size and molecular complexity of these compounds.

## Methods

### Compounds

We retrieved the chemical structure of 5500 small-molecules part of the Library of Integrate Network-based Cellular Signatures (LINCS - http://lincscloud.org) project in the form of SMILES string annotations (Supplementary Information). 4719 out of 5500 SMILES strings were retrieved according to their annotated ChemSpider ID (CSID) and PubChem ID (PID) in the NIH LINCS database. The remaining 779 NHS LINCS structures, for which no CSID or PID annotation was found, were retrieved by a web-API search in ChemSpider according to the molecule names. Six compounds were restricted structures. Thus, a final collection of 4927 LINCS unique structures was obtained. In addition, we retrieved chemical structures for the 1309 small-molecules part of the CMAP dataset (Connectivity Map).^2, 7^ 784 out of 1309 small-molecules were already present among the 4929 LINCS unique structures. Thus only 523 unique CMAP structures were retrieved as described before (**Supplementary Fig. 1**). The total number of chemical structure used for further analysis was thus equal to 5452.

The ChemAxon Standardizer tool (v. 14.9) was run to convert SMILES string annotations into 2D multi-SDF structural files.^74^ The “remove fragments” and “neutralize” options were used to fix all the molecular structures, to remove counter-ions and other various kinds of molecular fragments, which may be present in branded drug formulation but not useful in this work (e.g. besilates, mesilates, chlorides, bromides, sulphates, etc.). Protonation state of each structure was calculated with MoKa software v. 2.0 considering physiological pH 7.4.^75^

Finally, 3D minimized conformations were generated with the MMFF4x force-field in the MOE software (v. 2013)^76^ and stored as 3D multi-SDF structural files. The MMFF4x is the standard force-field parameterised for small organic molecules such as drugs. Partial charges are based on bond-charge increments. Conjugated nitrogens are considered as planar. Thus, a unique 3D multi-SDF file was obtained and used as input file for all the subsequent analyses.

### Physicochemical and pharmacokinetic properties

Starting from the 3D–coordinates multi-SDF file, each structure was the imported in the Volsurf+ v.1.5 software^27^ normalising their protonation state at pH 7.4. A set of 128 physicochemical and pharmacokinetic descriptors were calculated using Volsurf+ v. 1.5, using a grid spatial resolution of 0.5 Å. A final matrix of 5452 objects (drugs and chemical substances) and 128 descriptors was thus obtained. The molecular descriptors matrix was then visualised through the Principal Component Analysis (PCA) tool integrated in Volsurf+. Only the first five PCs were considered for the analysis. PCA score and loading plots are shown in **Supplementary Figure 2a** and **2b**. Analysis of the physicochemical descriptor distribution plots are shown in **Supplementary Figure 3**.

### 3D structural similarities by pharmacophore descriptors

The software FLAP v. 2.0^30^ was used to compute all-against-all pair-wise 3D structural similarities among the 5452 compounds. FLAP allows 3D molecular superimposition of two molecules and computes a pairwise similarity score based on Molecular Interaction Fields (MIFs), in order to evaluate type, strength, and direction of the interactions a molecule can have. The GRID tool,^23^ part of the FLAP software was used to compute the Molecular Interaction Fields based on three interaction probes: H, DRY and OH2. The hydrogen probe H is used to compute the shape of a small molecule. The hydrophobic probe DRY finds places at which hydrophobic atoms on the surface of a target molecule will make favourable interactions with hydrophobic ligand atoms. The probe OH2 represents polar and hydrophilic interactions mainly generated by hydrogen bond donor and acceptor functional groups and charges interactions. Four-point pharmacophores derived from the MIFs were used to align molecules with specific biological activity.^30, 77, 78^ The evaluation of MIF volume superimpositions between the two structures is reported as a similarity score ranging from 0 to 1 for each of the three probes. A global score (GLOB-Sum) is then obtained as the sum of the three scores of the individual probes. Higher GLOB-S values correspond to more similar structures. For this study, we transformed the GLOB-Sum similarity score matrix (**S**) of dimension 5452×5452 into a distance matrix defined as **D=1-S**/3.

Since the distance matrix is symmetric (i.e. the distance between A and B is the same as the distance between B and A), the total number of drug-pairs to consider is 14,859,426 (5452 × 5451 /2).

### Construction of the drug network

We ranked drug-pairs according to their structural distance in ascending order and considered as significant only those drug-pairs in the top 5% of the ranked list, as previously described by Iorio et. al.^4^ to reduce the total amount of egdes in the MANTRA network (The distance threshold is 0.51 when considering the 5452×5452 network or 0.65 when considering only the CMAP 1309x1309 sub-network). We then represented drugs as nodes connected by edges. The resulting Structural Drug Network has a giant connected component with 5312 nodes (i.e., drugs) out of 5,452 and 35,527 edges, corresponding to 5% of a fully connected network with the same number of nodes (14,859,426 edges) (**Supplementary Fig. 4**). In order to visualise and extract useful information from the SDN, we identified communities via the Affinity Propagation Clustering algorithm, as implemented in the R package apcluster (v. 1.3.5).^32, 79^ A community is defined as a group of nodes densely interconnected with each other and with fewer connections to nodes outside the group.^80^ Each community was coded with a numerical identifier, a colour, and one of its nodes was identified as the “exemplar” of the community, i.e., the drug whose effect best represents the effects of the other drugs in the community.^4^

### Validation of the Structural Drug Network

To validate the drug structural network, we assessed whether pairs of drugs connected by an edge in the network (i.e. structurally similar according to our distance) shared a common clinical application. To this end, we collected for each drug the correspondent Anatomical Therapeutic Chemical (ATC) code (version Index 2014). This drugs classification method developed by the World Health Organization in collaboration with the Drug Statistics Methodology (WHOCC),^81^ hierarchically classifies compounds according to five different levels: (1st level) Organ or system on which they act; (2nd level) Therapeutic class; (3rd level) Pharmacological subgroup; (4th level) Chemical subgroup; (5th level) Compound identifier. ATC code collisions often occur for the same drug. For instance, Aspirin has three distinct ATC codes: A01AD05 (drug for alimentary tract and metabolism), B01AC06 (blood agent as platelet inhibitor) and N02BA01 (nervous system agent as analgesic and antipyretic). In such cases we considered multiple ATC codes for the same drug in the network. ATC codes available from the WHOCC were 936 out of 5452 drugs (17%).

We then sorted drug-pairs according their structural distance in ascending order and for each drug-pair we checked whether they shared the same ATC to assess whether it was a True Positive (TP) or a False Positive (FP). **Supplementary Figure 6** reports the PPV=TP/(TP+FP) versus the drug-pair distance for different ATC code levels.

Furthermore, in order to assess whether a community in the drug network was enriched for a common ATC code, we counted the number of drugs with the same ATC code at the 4th level (pharmacological subclass) in community. We then computed a p-value for each community by applying the hypergeometric probability distribution test.

### Transcriptional Variability score

TV was computed for all the compounds having at least two profiles available in CMAP for the same cell line. The number of such small molecules for each cell line is: 1165 in MCF7, 398 in PC3, 32 in HL60, 2 in ssMCF7. We took advantage of the large majority of MCF7 experiments to avoid the problematic integration of TV values across different cell types and discarded all non-MCF7 data. About 15% of the CMAP small molecules have more than two profiles in MCF7 cells, producing an average of 16.08 “within-molecule” profile pairs and a maximum of 630 (for tanespimycin). To obtain the TV for a small molecule we computed the median of all the distances between such pairs. The pairwise distance is based on the enrichment of the top (bottom) genes of one profile among the top (bottom genes) of the other profile and vice-versa, as detailed in Iorio et. al.^4^ Since the TV is based on the same transcriptional distance measure used to derive the transcriptional network in Iorio et al^4^, we set as a significance threshold for the TV the same threshold used to derive the transcriptional network (TV_th_=0.8).

### Phospholipidosis stress signature

The PLD stress signature was built by merging together 35 PRLs (prototype ranked lists), corresponding to drugs searched in the literature known to induce PLD,^45^ into a single node using the Kruskal Algorithm strategy and the Borda Merging Method implemented the online tool MANTRA (http://mantra.tigem.it)and previously described ^3^. Briefly, the algorithm first searches for the two ranked lists with the smallest Spearman’s Footrule distance. Then it merges them using the Borda Merging Method, obtaining a new ranked list of genes. The process restarts until only one list remains.

### HCS (High Content Screening) assays

*TFEB nuclear translocation:* To quantify TFEB subcellular localization, a high-content assay upon the compound treatments indicated was performed using stable HeLa cells overexpressing TFEB-GFP according to our previous protocols (Medina et al, 2015). *Lysosome, Golgi and Endoplasmic Reticulum assays*: HeLa cells were seeded in a 384-well plate, incubated for 24h and treated with the different compounds at 0.1, 1 and 10 μM for additional 24h. After that cells were fixed with 4% paraformaldehyde (for LAMP1 and GM130 stainings) or ice-cold methanol (for PDI staining) and permeabilized/blocked with 0.05% (w/v) saponin, 0.5% (w/v) BSA and 50 mM NH4Cl in PBS (blocking buffer). LAMP-1, GM130 and PDI detection was performed by incubating with the corresponding primary antibodies (anti-LAMP1, Santa Cruz Biotechnology; anti-GM130 and anti-PDI, Cell Signaling Technology) followed by the incubation with an AlexaFluor-conjugated secondary antibodies (Life Technologies) diluted in blocking buffer. LysoTracker Red DND-99 (Life Technologies) staining was performed by the incubating the dye for the last 30 minutes before fixation. DAPI and CellMask Deep Red Plasma membrane Stain (Life Technologies) were used for nuclei and plasma membrane staining, respectively. Images of of lysosomes (LAMP-1 and LysoTracker Red DND-99), Golgi (GM130) and ER (PDI) were acquired using an automated confocal microscopy (Opera High Content System, Perkin-Elmer). The fluorescent intensity and area of the different stainings were analyzed by using dedicated scripts developed in the Columbus Image Data Management and Analysis Software (Perkin-Elmer).

*High Content Lipid accumulation assay:* LipidTOX green phospholipidosis detection reagent (Life Technologies) was added to the cells along with the different compounds at the indicated concentrations for 48h before fixation with 4% paraformaldehyde. DAPI and CellMask Deep Red Plasma membrane Stain (Life Technologies) were used for nuclei and plasma membrane staining, respectively. Lysosomal phospholipid accumulation was analyzed by measuring fluorescent dye intensity using an automated confocal microscopy (Opera High Content System, Perkin-Elmer) and a Columbus Image Data Management and Analysis Software (Perkin-Elmer).

## Acknowledgments

The authors are grateful to the Bioinformatics Core (TIGEM). This work was supported by a Fondazione Telethon Grant (TGM11SB1) to DdB.

## Competitive financial interests

The authors declare no competing financial interests.

## Materials and Correspondence

Author: Diego di Bernardo

## References

1 Verbist, B. et al. Using transcriptomics to guide lead optimization in drug discovery projects: Lessons learned from the QSTAR project. Drug Discov Today 20, 505–513 (2015).

2 Lamb, J. et al. The Connectivity Map: Using Gene-Expression Signatures to Connect Small Molecules, Genes, and Disease. Science 313, 1929–1935 (2006).

3 Cheng, J., Yang, L., Kumar, V. & Agarwal, P. Systematic evaluation of connectivity map for disease indications. Genome Medicine 6, 95 (2014).

4 Iorio, F. et al. Discovery of drug mode of action and drug repositioning from transcriptional responses. Proceedings of the National Academy of Sciences 107, 14621–14626 (2010).

5 Woo, J.H. et al. Elucidating Compound Mechanism of Action by Network Perturbation Analysis. Cell 162, 441–451 (2015).

6 Kidd, B.A. et al. Mapping the effects of drugs on the immune system. Nat Biotechnol 34, 47–54 (2016).

7 Lamb, J. The Connectivity Map: a new tool for biomedical research. Nat Rev Cancer 7, 54–60 (2007).

8 Iorio, F., Rittman, T., Ge, H., Menden, M. & Saez-Rodriguez, J. Transcriptional data: a new gateway to drug repositioning? Drug Discov Today 18, 350–357 (2013).

9 Bajorath, J. et al. Navigating structure–activity landscapes. Drug Discovery Today 14, 698–705 (2009).

10 Geppert, H., Vogt, M. & Bajorath, J. Current trends in ligand-based virtual screening: molecular representations, data mining methods, new application areas, and performance evaluation. J Chem Inf Model 50, 205–216 (2010).

11 Heikamp, K. & Bajorath, J. The Future of Virtual Compound Screening. Chemical Biology & Drug Design 81, 33–40 (2013).

12 Shim, J. & Mackerell, A.D., Jr. Computational ligand-based rational design: Role of conformational sampling and force fields in model development. Medchemcomm 2, 356–370 (2011).

13 Sirci, F. et al. Virtual fragment screening: discovery of histamine H3 receptor ligands using ligand-based and protein-based molecular fingerprints. J Chem Inf Model 52, 3308–3324 (2012).

14 Stumpfe, D. & Bajorath, J. Activity Cliff Networks for Medicinal Chemistry. Drug Development Research 75, 291–298 (2014).

15 Vogt, M. & Bajorath, J. Chemoinformatics: A view of the field and current trends in method development. Bioorganic & Medicinal Chemistry 20, 5317–5323 (2012).

16 Backman, T.W., Cao, Y. & Girke, T. ChemMine tools: an online service for analyzing and clustering small molecules. Nucleic Acids Res 39, W486–491 (2011).

17 Ma, X.H. et al. Comparative analysis of machine learning methods in ligand-based virtual screening of large compound libraries. Combinatorial chemistry & high throughput screening 12, 344–357 (2009).

18 Ravindranath, A.C. et al. Connecting gene expression data from connectivity map and in silico target predictions for small molecule mechanism-of-action analysis. Mol Biosyst 11, 86–96 (2015).

19 Khan, S.A. et al. Identification of structural features in chemicals associated with cancer drug response: a systematic data-driven analysis. Bioinformatics 30, 1497–504 (2014).

20 Iskar, M. et al. Drug-induced regulation of target expression. PLoS Comput Biol 6 (2010).

21 Hizukuri, Y., Sawada, R. & Yamanishi, Y. Predicting target proteins for drug candidate compounds based on drug-induced gene expression data in a chemical structure-independent manner. BMC Med Genomics 8, 82 (2015).

22 Carosati, E., Sciabola, S. & Cruciani, G. Hydrogen Bonding Interactions of Covalently Bonded Fluorine Atoms:L From Crystallographic Data to a New Angular Function in the GRID Force Field. Journal of Medicinal Chemistry 47, 5114–5125 (2004).

23 Goodford, P.J. A computational procedure for determining energetically favorable binding sites on biologically important macromolecules. Journal of Medicinal Chemistry 28, 849–857 (1985).

24 Carrella, D. et al. Mantra 2.0: an online collaborative resource for drug mode of action and repurposing by network analysis. Bioinformatics 30, 1787–1788 (2014).

25 Iorio, F., Isacchi, A., di Bernardo, D. & Brunetti-Pierri, N. Identification of small molecules enhancing autophagic function from drug network analysis. Autophagy 6, 1204–1205 (2010).

26 Cruciani, G., Crivori, P., Carrupt, P.A. & Testa, B. Molecular fields in quantitative structure–permeation relationships: the VolSurf approach. Journal of Molecular Structure: THEOCHEM 503, 17–30 (2000).

27 Cruciani, G., Pastor, M. & Guba, W. VolSurf: a new tool for the pharmacokinetic optimization of lead compounds. European Journal of Pharmaceutical Sciences 11, Supplement 2, S29–S39 (2000).

28 Lipinski, C.A. Lead- and drug-like compounds: the rule-of-five revolution. Drug Discov Today Technol 1, 337–341 (2004).

29 Lipinski, C.A., Lombardo, F., Dominy, B.W. & Feeney, P.J. Experimental and computational approaches to estimate solubility and permeability in drug discovery and development settings. Advanced Drug Delivery Reviews 23, 3–25 (1997).

30 Baroni, M., Cruciani, G., Sciabola, S., Perruccio, F. & Mason, J.S. A common reference framework for analyzing/comparing proteins and ligands. Fingerprints for Ligands and Proteins (FLAP): theory and application. J Chem Inf Model 47, 279–294 (2007).

31 Bodenhofer, U., Kothmeier, A. & Hochreiter, S. APCluster: an R package for affinity propagation clustering. Bioinformatics 27, 2463–2464 (2011).

32 Frey, B.J. & Dueck, D. Clustering by passing messages between data points. Science 315, 972–976 (2007).

33 Baliga, B.S., Pronczuk, A.W. & Munro, H.N. Mechanism of cycloheximide inhibition of protein synthesis in a cell-free system prepared from rat liver. J Biol Chem 244, 4480–4489 (1969).

34 Jimenez, A., Carrasco, L. & Vazquez, D. Enzymic and nonenzymic translocation by yeast polysomes. Site of action of a number of inhibitors. Biochemistry 16, 4727–4730 (1977).

35 McKeehan, W. & Hardesty, B. The mechanism of cycloheximide inhibition of protein synthesis in rabbit reticulocytes. Biochemical and Biophysical Research Communications 36, 625–630 (1969).

36 Nadanaciva, S. et al. A high content screening assay for identifying lysosomotropic compounds. Toxicol In Vitro 25, 715–723 (2011).

37 Petersen, Nikolaj H.T. et al.

38 Ellegaard, A.-M. et al.

39 Roy, M., Dumaine, R. & Brown, A.M. HERG, a primary human ventricular target of the nonsedating antihistamine terfenadine. Circulation 94, 817–823 (1996).

40 Zhou, Z., Vorperian, V.R., Gong, Q., Zhang, S. & January, C.T. Block of HERG Potassium Channels by the Antihistamine Astemizole and its Metabolites Desmethylastemizole and Norastemizole. Journal of Cardiovascular Electrophysiology 10, 836–843 (1999).

41 Morissette, G., Lodge, R. & Marceau, F. Intense pseudotransport of a cationic drug mediated by vacuolar ATPase: procainamide-induced autophagic cell vacuolization. Toxicol Appl Pharmacol 228, 364–377 (2008).

42 Ashoor, R., Yafawi, R., Jessen, B. & Lu, S. The Contribution of Lysosomotropism to Autophagy Perturbation. PLoS One 8, e82481 (2013).

43 Kazmi, F. et al. Lysosomal Sequestration (Trapping) of Lipophilic Amine (Cationic Amphiphilic) Drugs in Immortalized Human Hepatocytes (Fa2N-4 Cells). Drug Metabolism and Disposition 41, 897–905 (2013).

44 Marceau, F. et al. Cation trapping by cellular acidic compartments: beyond the concept of lysosomotropic drugs. Toxicol Appl Pharmacol 259, 1–12 (2012).

45 Muehlbacher, M., Tripal, P., Roas, F. & Kornhuber, J. Identification of Drugs Inducing Phospholipidosis by Novel in vitro Data. Chemmedchem 7, 1925–1934 (2012).

46 Halliwell, W.H. Cationic amphiphilic drug-induced phospholipidosis. Toxicol Pathol 25, 5360 (1997).

47 Goracci, L., Ceccarelli, M., Bonelli, D. & Cruciani, G. Modeling Phospholipidosis Induction: Reliability and Warnings. Journal of Chemical Information and Modeling 53, 1436–1446 (2013).

48 Sun, H. et al. Are hERG channel blockers also phospholipidosis inducers? Bioorganic & medicinal chemistry letters 23, 4587–4590 (2013).

49 Anderson, N. & Borlak, J. Drug-induced phospholipidosis. FEBS Letters 580, 5533–5540 (2006).

50 Lu, S., Sung, T., Lin, N., Abraham, R.T. & Jessen, B.A. Lysosomal adaptation: How cells respond to lysosomotropic compounds. PLOS ONE 12, e0173771 (2017).

51 Napolitano, F., Sirci, F., Carrella, D. & di Bernardo, D. Drug-set enrichment analysis: a novel tool to investigate drug mode of action. Bioinformatics (2015).

52 Napolitano, G. & Ballabio, A. TFEB at a glance. Journal of cell science 129, 2475–2481 (2016).

53 Martina, J.A., Chen, Y., Gucek, M. & Puertollano, R. MTORC1 functions as a transcriptional regulator of autophagy by preventing nuclear transport of TFEB. Autophagy 8, 903–914 (2012).

54 Roczniak-Ferguson, A. et al. The transcription factor TFEB links mTORC1 signaling to transcriptional control of lysosome homeostasis. Science signaling 5, ra42 (2012).

55 Settembre, C. et al. A lysosome-to-nucleus signalling mechanism senses and regulates the lysosome via mTOR and TFEB. The EMBO journal 31, 1095–1108 (2012).

56 Sardiello, M. et al. A gene network regulating lysosomal biogenesis and function. Science 325, 473–477 (2009).

57 Settembre, C. et al. TFEB links autophagy to lysosomal biogenesis. Science 332, 1429–1433 (2011).

58 Medina, Diego L. et al. Transcriptional Activation of Lysosomal Exocytosis Promotes Cellular Clearance. Developmental Cell 21, 421–430 (2011).

59 Medina, D.L. et al. Lysosomal calcium signalling regulates autophagy through calcineurin and TFEB. Nature cell biology 17, 288–299 (2015).

60 Carrella, D. et al. Computational drugs repositioning identifies inhibitors of oncogenic PI3K/AKT/P70S6K-dependent pathways among FDA-approved compounds. Oncotarget (2016).

61 Jin, Y. et al. Antineoplastic mechanisms of niclosamide in acute myelogenous leukemia stem cells: inactivation of the NF-kappaB pathway and generation of reactive oxygen species. Cancer Res 70, 2516–2527 (2010).

62 Ishii, I., Harada, Y. & Kasahara, T. Reprofiling a classical anthelmintic, pyrvinium pamoate, as an anti-cancer drug targeting mitochondrial respiration. Frontiers in oncology 2, 137 (2012).

63 Fonseca, B.D. et al. Structure-activity analysis of niclosamide reveals potential role for cytoplasmic pH in control of mammalian target of rapamycin complex 1 (mTORC1) signaling. J Biol Chem 287, 17530–17545 (2012).

64 Newman, R.A., Yang, P., Pawlus, A.D. & Block, K.I. Cardiac glycosides as novel cancer therapeutic agents. Molecular interventions 8, 36–49 (2008).

65 Wang, Y.C., Chen, S.L., Deng, N.Y. & Wang, Y. Network predicting drug’s anatomical therapeutic chemical code. Bioinformatics 29, 1317–1324 (2013).

66 Krishnan, A.V., Swami, S. & Feldman, D. Estradiol inhibits glucocorticoid receptor expression and induces glucocorticoid resistance in MCF-7 human breast cancer cells. J Steroid Biochem Mol Biol 77, 29–37 (2001).

67 Zhang, Y., Leung, D.Y.M., Nordeen, S.K. & Goleva, E. Estrogen Inhibits Glucocorticoid Action via Protein Phosphatase 5 (PP5)-mediated Glucocorticoid Receptor Dephosphorylation. The Journal of Biological Chemistry 284, 24542–24552 (2009).

68 Carollo, M., Parente, L. & D’Alessandro, N. Dexamethasone-induced cytotoxic activity and drug resistance effects in androgen-independent prostate tumor PC-3 cells are mediated by lipocortin 1. Oncol Res 10, 245–254 (1998).

69 Zhang, C. et al. Corticosteroid-induced chemotherapy resistance in urological cancers. Cancer Biol Ther 5, 59–64 (2006).

70 Hamid, N. & Krise, J.P. in Lysosomes: Biology, Diseases, and Therapeutics 423–444 (John Wiley & Sons, Inc., 2016).

71 Liu, J., Lee, J., Hernandez, M.A.S., Mazitschek, R. & Ozcan, U. Treatment of Obesity with Celastrol. Cell 161, 999–1011 (2015).

72 Chen, B. & Butte, A.J. Leveraging big data to transform target selection and drug discovery. Clinical pharmacology and therapeutics 99, 285–297 (2016).

73 Sirota, M. et al. Discovery and preclinical validation of drug indications using compendia of public gene expression data. Science translational medicine 3, 96ra77 (2011).

74 JChem 14.9.15, 2014, ChemAxon (http://www.chemaxon.com)”

75 Milletti, F., Storchi, L., Sforna, G. & Cruciani, G. New and Original pKa Prediction Method Using Grid Molecular Interaction Fields. Journal of Chemical Information and Modeling 47, 2172–2181 (2007).

76 Molecular Operating Environment (MOE), 2013.08; Chemical Computing Group Inc., 1010 Sherbooke St. West, Suite #910, Montreal, QC, Canada, H3A 2R7, 2016.

77 Cross, S., Baroni, M., Carosati, E., Benedetti, P. & Clementi, S. FLAP: GRID Molecular Interaction Fields in Virtual Screening. Validation using the DUD Data Set. Journal of Chemical Information and Modeling 50, 1442–1450 (2010).

78 Cross, S. & Cruciani, G. Grid-derived structure-based 3D pharmacophores and their performance compared to docking. Drug Discovery Today: Technologies 7, e213–e219 (2010).

79 De Baets, B. & Mesiar, R. Metrics and T-Equalities. Journal of Mathematical Analysis and Applications 267, 531–547 (2002).

80 Newman, M.E.J. Modularity and community structure in networks. Proceedings of the National Academy of Sciences 103, 8577–8582 (2006).

81 WHO Collaborating Centre for Drug Statistics Methodology, ATC classification index with DDDs, 2014. Oslo 2014.

